# Mapping the peripheral immune landscape of Parkinson’s disease patients with single-cell sequencing

**DOI:** 10.1101/2024.07.26.605020

**Authors:** Gael Moquin-Beaudry, Lovatiana Andriamboavonjy, Sebastien Audet, Laura K Hamilton, Antoine Duquette, Sylvain Chouinard, Michel Panisset, Martine Tetreault

**Affiliations:** University of Montreal Hospital Research Center (CRCHUM), Montreal H2X 0A9, Canada; Department of Neuroscience, University of Montreal, Montreal H3T 1J4, Canada; Department of Medicine, University of Montreal, Montreal H3T 1J4, Canada; André-Barbeau movement disorders unit, University of Montreal Hospital (CHUM), Montreal H2X 3E4, Canada; Neurology service, University of Montreal Hospital (CHUM), Montreal H2X 3E4, Canada

**Author notes:** Correspondence to: Martine Tetreault, Department of Neuroscience, University of Montreal, CRCHUM-Tour Viger 900, Saint-Denis Street, R09.706, Montreal, QC H2X 0A9, Canada. Contributed equally (Gael Moquin-Beaudry, Lovatiana Andriamboavonjy, Sebastien Audet).

**Keywords:** Stress response, inflammation, monocyte activation, TCR sequencing, activation signature

## Abstract

Parkinson’s Disease (PD) is most recognized for its impact on the central nervous system. However, recent breakthroughs underscore the crucial role of interactions between central and peripheral systems in PD’s pathogenesis. The spotlight is now shifting as we explore beyond the central nervous system, discovering that peripheral changes such as inflammatory dysfunctions may predict the rate of disease progression and severity. Despite more than 200 years of research on PD, reliable diagnostic or progression biomarkers and effective disease-modifying treatments are still lacking. Additionally, the cellular mechanisms that drive changes in immunity are largely unknown. Thus, understanding peripheral immune signatures could lead to earlier diagnosis and more effective treatments for PD. Here, we sought to define the transcriptomic alterations of the complete peripheral immune cell compartment by single-cell RNA- and T-cell receptor-sequencing with hopes of uncovering PD signatures and potential peripheral blood biomarkers.

Following transcriptional profiling of 78 876 cells from 10 healthy controls and 14 PD donors, we observed five major classes of immune cell types; myeloid (monocytes, dendritic cells) and lymphoid (T, B, natural killer) cells from which we identified 38 cellular subtypes following bioinformatic re-clustering. Comparing immune cell subtypes and phenotypes between PD patients and healthy controls revealed notable features of PD: 1) a significant shift of classical CD14^+^ monocytes towards an activated CD14^+^/CD83^+^ state, 2) changes in lymphocyte subtype abundance, including a significant decrease in CD4^+^ naive and mucosal-associated invariant T-cells subtypes, along with an increase in CD56^+^ natural killer cells, 3) the identification by T-cell receptor sequencing of several PD specific T-cell clones shared between multiple patients, suggesting the implication of common epitopes in PD pathogenesis, 4) a notable increase in the expression of activation signature genes, including the AP-1 stress-response transcription factor complex, across all PD cell types. This signal was not present in atypical parkinsonism patients with multiple systems atrophy or progressive supranuclear palsy.

Overall, we present a comprehensive atlas of peripheral blood mononuclear cells from control and PD patients which should serve as a tool to improve our understanding of the role the immune cell landscape plays in PD pathogenesis.

## Introduction

Parkinson’s Disease (PD) is a complex neurodegenerative disorder characterized by the progressive loss of dopaminergic neurons in the substantia nigra pars compacta, and the intracytoplasmic neuronal accumulation of the α-synuclein protein.^1,2,3^ PD currently affects over 10 million people worldwide, with an increasing global prevalence estimated at 17 million by 2040.^4,5^ Unfortunately, there is no biomarker for PD to date, and misdiagnosis occurs in up to 16% of cases.^6^ Thus, these misdiagnosed patients are deprived of necessary care, which impacts their disease prognosis.^7^ Moreover, there is no cure for PD and the available therapies only temporarily ease patients’ symptoms.^8^ Even though they are still under investigation, disease-modifying treatments are assumed to maximize benefits if introduced at an early-stage of the disease, before the apparition of the motor symptoms.^9,10,11^ Given the widespread peripheral changes observed early in PD, using easily accessible peripheral tissues for biomarker discovery is highly promising. Indeed, the involvement of multisystem inflammation in PD pathophysiology has been suspected for some time with the first report dating back to 1988 when McGeer *et al.*^12^ noted microglial activation in the substantia nigra of PD postmortem brains. The inflammation hypothesis has been reinforced by the identification of pro-inflammatory cytokines in the brain tissue and cerebrospinal fluid of PD patients.^13,14^ Subsequent studies have observed the infiltration of peripheral immune cells into the brain, believed to breach the blood-brain barrier and amplify the microglial reaction.^15,16^ Of note, these immune alterations are also present in the periphery, showing elevated levels of proinflammatory cytokines in serum and an altered blood immune cell profile.^17^ This connection between the central and peripheral immune system, along with the alterations seen in both, offers an opportunity to discover blood-based biomarkers for the diagnosis and prognosis of PD.

The rise of single-cell technologies has allowed for the characterization of complex systems at single-cell resolution, providing a novel unbiased way to identify gene signatures and functional alterations simultaneously across the entire cellular landscape. In recent years, multiple groups have applied such technologies to accelerate our understanding of PD pathogenesis in the brain.^18,19^ However, no PD studies have interrogated the complete peripheral immune compartment in this fashion.^20,21^ We believe that by studying these peripheral changes, we can enhance early diagnosis and help identify new therapeutic targets for PD.

In this study, we investigated the complete peripheral blood mononuclear cell compartment (PBMCs), encompassing myeloid (monocytes, dendritic cells (DC) and lymphoid (T, B, natural killer (NK) cells). These cells are ideal for investigation due to their pivotal roles in immune response, metabolism, and intercellular communication, as well as their ease of collection from live subjects. By employing single-cell RNA (sc-RNAseq) and T-cell receptor (TCR) sequencing simultaneously, our aim was to uncover precise immune-cell signatures unique to PD. Comparing the immune cell profiles between PD patients and healthy controls successfully unveiled significant differences indicative of distinct immune system changes in PD. Overall, our study provides a comprehensive atlas of PBMCs in both healthy individuals and PD patients. This resource enhances our understanding of the immune cell landscape in PD and may lead to novel insights into disease mechanisms and potential therapeutic targets.

## Materials and methods

### Human PBMCs

Blood samples were obtained from patients with sporadic PD, multiple system atrophy (MSA) and progressive supranuclear palsy (PSP), as well as age and sex matched healthy donors after informed consent was obtained in accordance with the institutional Ethics committee protocol #20.367 in the context of a biobank. Samples were recovered in EDTA vacutainer tubes and processed within 4 hours of harvest. Blood was diluted with equal volume of PBS-2mM EDTA, deposited over 12mL of Ficoll-Paque (GE Healthcare) and centrifuged as per manufacturer’s instructions. Buffy coats were harvested, washed with PBS-2mM EDTA and aliquoted with 10% DMSO (Sigma-Aldrich) in fetal bovine serum (FBS, Gibco) freezing solution for long term storage in liquid nitrogen.

### Single-cell processing

Donor cells were distributed evenly between 5-plex batches to limit batch artifacts. Cells were processed on ice and centrifugations operated at 4°C when possible.

#### Thawing

Samples were quickly thawed in a 37°C water bath until only a small bit of ice remained in the cryovial and transferred into precooled 15mL Falcon tubes containing 40µL nuclease S7 (Sigma Aldrich) which was quenched after 30 seconds by slowly adding 5mL pure FBS followed by centrifugation at 4°C, 200 RCF for 10 minutes. The resulting pellet was washed twice with 5mL PBS containing 0.5% bovine serum albumin and 2mM EDTA, resuspended in 10% FBS RPMI at approximately 2×10⁶ cells/mL, and allowed to rest for 2 hours in a 37°C incubator before pursuing.

#### Dead cell removal

Approximately 2×10⁶ cells were treated with a dead cell removal kit (Miltenyi) as per manufacturer’s protocol. Briefly, cells were harvested and washed after centrifugation and resuspended in 30µL of Dead Cell Removal Microbeads. After a 15 minute incubation at room temperature, 470µL of freshly prepared binding buffer was added and loaded onto pre-rinsed MACS MS columns (Miltenyi). Flowthrough was collected and columns rinsed four times, harvesting the flowthrough. Cells were centrifuged and resuspended in 1mL of 10% FBS RPMI in a precooled 1.5 mL LoBind tube (Eppendorf) and kept on ice.

#### Cell hashing and pooling

Cells were centrifuged and resuspended in 100µL of Fc blocking solution (Biolegend). After a 10 minute incubation on ice, 0.5µg of the appropriate TotalSeq-B antibody (Biolegend) was added to each sample and incubated for 20 minutes on ice. Cells were washed in filtered PBS +2% BSA +0.01% Tween 20 (Sigma Aldrich) three times and finally resuspended in 250µL PBS. Cells were counted manually via hemocytometer and 2×10⁵ cells for each suspension were pooled into their designated 5-plex run mix (10⁶ cells per mix), passed through 40µm cell strainer (Miltenyi), rinsed with PBS, centrifuged, and resuspended in 300µL of PBS. Cell mixes were finally counted manually via hemocytometer to confirm the loading volume, viability, and single cellularity.

#### 10X Genomics

5-plex mixes were processed using 10X Genomics 5’ v1.1 kit, overloading the Chromium Next GEM Chip G lane with 7.2×10^4^ cells for an expected yield of 3×10^4^ recovered cells per mix. Following droplet generation, gene expression, V(D)J, and feature barcoding libraries were generated using the manufacturer’s protocol.

### Next generation sequencing

Libraries were sequenced on a NovaSeq 6000 S4 (Illumina) using the 10X recommended read parameters for 5ʹ v.1.1 libraries: paired-end, single indexing, Read 1: 26 bp; Read 2: 91 bp; i7: 8 bp. Libraries were multiplexed with a targeted sequencing depth of 50,000 reads/cell for GEX libraries and 5,000 reads/cell for V(D)J and feature barcoding libraries. Sequencing was performed at the CHU de Québec-Université Laval’s genomic center.

### Single-cell data preprocessing

All data preprocessing was performed on Digital Research Alliance of Canada’s high performance computing clusters Béluga and Narval. Count matrices were generated using Cellranger v6.0.1 (10X Genomics) with the GRCh38 genome.^22^ GEX and feature barcoding libraries were processed using the “count” function and V(D)J libraries with the “vdj” function.

### Single-cell analysis tools

Single-cell analysis was performed in R (v4.3.1) mainly using the single-cell analysis package Seurat v4.3.0.1. Some figures were generated using ggplot2 v3.4.2, scCustomize v1.1.3, dittoSeq v1.12.2 and ggpubr v0.6.0.^23,24,25,26,27^ Other R packages were employed as described in the relevant methods section.

### Sample demultiplexing

For each 5-plex sample, count matrices were imported into R and subsetted to segregate feature barcoding and gene expression counts into distinct assays within the Seurat object. Donor demultiplexing of 5-plex barcodes was done by associating hashtag oligonucleotides (HTO) to genotypes. HTO calls were obtained from antibody feature barcoding using Seurat’s “HTODemux” function, while the genotype demultiplexing tool souporcell v2.0.40 was used to obtain genotype calls.^28,29^ Once genotypes were attributed to each donor, genotype demultiplexing was used to attribute cells to each donor due to lower dropout rate than the HTO method. Doublets were filtered out.

Demultiplexed samples were subsequently annotated with their respective donor’s metadata. Downstream integration was conducted using Seurat’s integration pipeline with default settings and using the “SCTransform” normalization method.^30,31^ Of note, ribosomal (^RPL* and ^RPS*) and mitochondrial (^MT*) genes were removed from the 5,000 integration features identified by SelectIntegrationFeatures function.

### Initial quality control and clustering

Standard cell quality control was run and cells with high or low feature count (<700 & >5000) and high mitochondrial content (>10%) were filtered out. Principal Component Analysis and subsequent Uniform Manifold Approximation and Projection (UMAP) calculations were done using previously filtered integration features, to which T-cell receptor (^TR[ABDG][VDJ]*) and Immunoglobulin (^IG[KHL]V*) genes were additionally filtered out. Jackstraw plots were employed to assess the significance of principal components, aiding in the determination of the appropriate number of dimensions to use for clustering.^32^

### Cell annotation and filtering

A first automatic cell annotation was performed using Seurat’s reference mapping function to project our query dataset on the reference dataset’s UMAP projection, transfer cell type annotations (predicted.celltype.l1 and predicted.celltype.l2) and impute protein expression from the reference’s CITE-seq data. The reference dataset used in this case was Hao *et al*.’s^33^ Human PBMC (162k cells with 228 antibodies). Low granularity cell type prediction score was used as a cell quality filtering metric with cells having a calling score below 0.85 being filtered out. *De novo* UMAP calculation was performed, and low confidence cells were filtered out. This low confidence label was manually attributed for multiple reasons, such as major discrepancies between cell type annotation and location in the UMAP, heterogeneous or hybrid subclusters, and major discrepancies between high- and low-level cell type annotation. A second automatic cell type annotation procedure was run using the SciBetR v0.1.0 package with Stephenson *et al*.’s ^34^ COVID-19 PBMC Ncl-Cambridge-UCL reference dataset for a richer annotation.^35^ Cell type annotations were further manually curated for clarity.

### Differential gene expression

Differential expression (DE) analysis was performed using Seurat’s “FindMarker” function with standard parameters on library size normalized UMI counts to identify DE genes between conditions within broad cell types. DE genes were defined as having an adjusted P value <0.05. Gene module score was calculated by Seurat’s “AddModuleScore” function with the genes of interest.

### Population composition analysis

Determination of differential cell population abundance between conditions was computed using the DCATS beta-binomial generalized linear statistical model to compare annotated clusters.^36^ The miloR method of differential abundance analysis was also applied to validate differences between PD and control (CTRL) donor samples (Supplementary Fig. 1).^37^

### TCR sequencing and analysis

Output from CellRanger’s “vdj” function with reference = refdata-celranger-vdj-GRCh38-alts-ensembl-5.0.0 was analyzed using the R package scRepertoire v1.3.5.^38^ Clonotypes were defined as cells sharing the same VDJC genes and complementarity-determining region 3 (CDR3) nucleotide for TCRs ɑ and β. The tool also allows for easy visualization and integration of TCR data into the Seurat object.

To identify common antigen-specific TCRs between donors based on CDR3 sequence similarity, βCDR3 sequences were clustered using the GIANA tool: Geometry Isometry based TCR AligNment Algorithm v4.1 python software.^39^ The isometric encoding of amino acid CDR3 sequences is used to define clusters by filtering out unique sequences using approximate nearest neighbor search and identifying clusters using pairwise comparison with the remaining sequences. Data shown was obtained using a threshold for Smith-Waterman alignment score (-S option) of 3.5 and Isometric distance threshold (-t option) of 10. Resulting sequence clusters were sorted by donor and by condition to identify clonotypes shared between multiple individuals and more precisely only among PD patients. Cluster sequences visualization was done using the ggseqlogo v0.1 R package.^40^

### Single-cell trajectory analysis

Relevant clusters were subset from the Seurat object and directly transformed into a “cell_data_set” object using SeuratWrapper v0.3.1.^41^ Subsequent processing was performed with the Monocle3 v1.3.3 R package.^42,43^ Cells were partitioned and re-clustered through the UMAP while the defined resolution was chosen to be comparable to previous clustering. Gene pseudotime trajectory inference was calculated using classical CD14^+^ monocytes as the starting state. Subsequent annotations utilized gene modules that most strongly correlated with each cluster while significantly driving trajectory signals (*Q* < 0.05).

### Statistics

Statistical analysis of data was performed mainly using the base R and Seurat R packages. The number of donors was not predetermined by a statistical method, but sample size is comparable or higher than those commonly used in similar single-cell literature.^20^ Experiments used mostly non-blinded analyses. Experimental collection and processing of samples were performed non-blinded to ensure an even distribution of samples by age, sex, and condition of donors and used automatic and field standard analysis procedures requiring donor-dependent metadata to be performed.

Exclusion of data points from this analysis occurred when cells were labeled as low-quality or defined as doublets by demultiplexing tools (see previous methods sections). All filtering was performed using field standard techniques and thresholds, and all filtered data points are present in raw deposited files.

## Results

### Comprehensive Single-Cell Analysis of Peripheral Immune Cells in PD

Gaining detailed insight into PBMC subpopulations is challenging due to their heterogeneous composition, dynamic nature and diverse functional roles in the immune response. To identify cellular or molecular biomarkers for tracking and diagnosing PD, we generated a comprehensive high-quality sc-RNAseq dataset of PBMCs from 14 PD patients and 10 age- and sex-matched healthy controls (CTRL) (Fig. 1A; Supplementary Table 1). Sample multiplexing was employed to maximize efficiency, and the samples from a single batch were demultiplexed using hashtag oligo-bound antibodies and genotypic computational approaches. This enabled the identification of individual donor cells and the removal of dual-donor doublets. Data integration was performed using the R package Seurat to mitigate potential batch effects from sample processing (see Methods). Following doublet removal and cell quality control, we obtained 78,876 cells, with a median of 4,349 unique molecular identifiers (UMIs) and 1,606 features per cell. T-cell receptor (TCR) data was also obtained for 57,745 of these cells.

**Figure 1.**
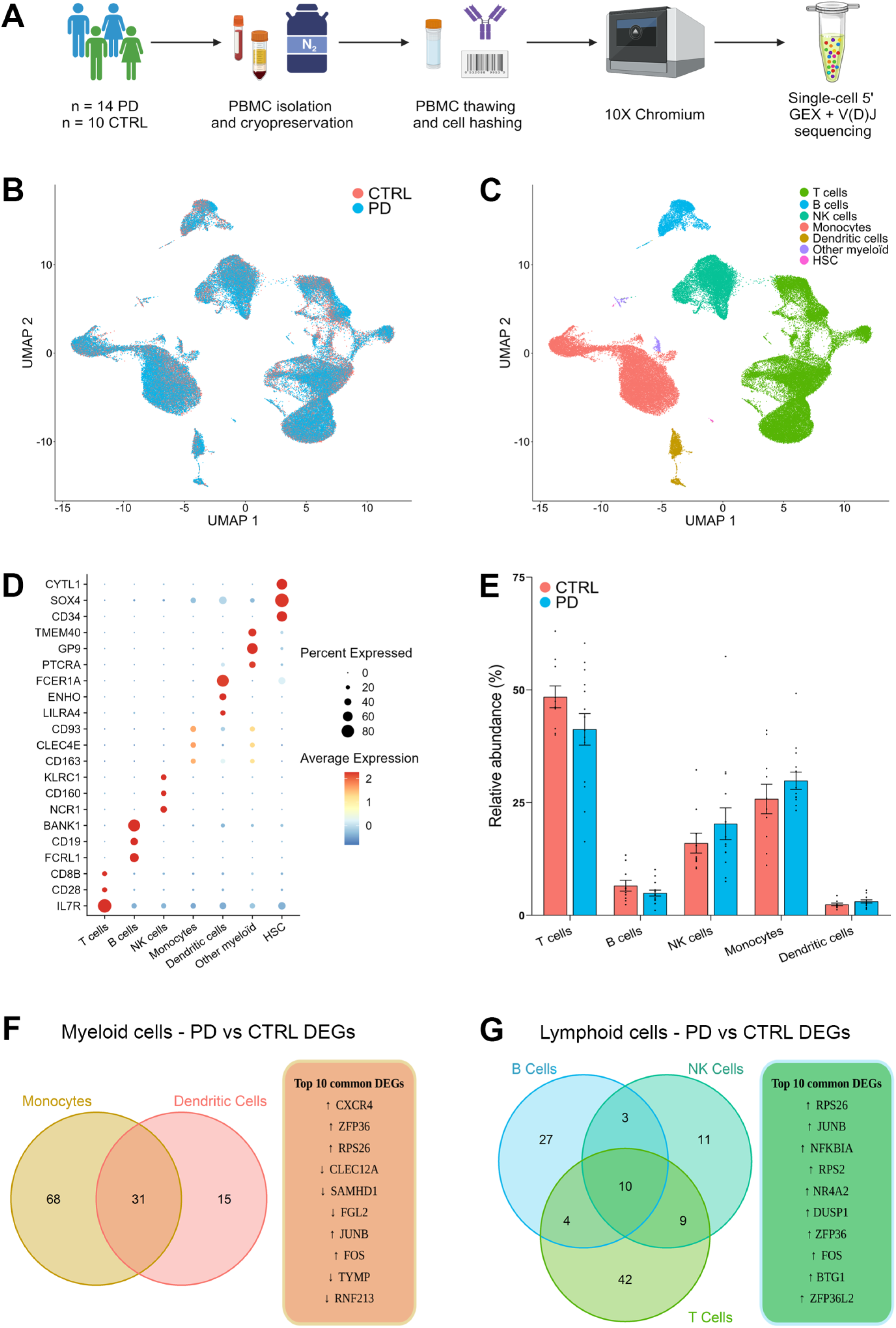
Comprehensive Single-Cell Analysis of Peripheral Immune Cells in PD. **(A)** Schematic of sample origin and processing. Cryopreserved peripheral blood mononuclear cells (PBMCs) from 14 Parkinson’s Disease (PD) patients and 10 healthy controls (CTRL) were prepared, multiplexed, and processed using 10X Genomics 5’ v1.1 chemistry to obtain gene expression (GEX) and V(D)J libraries. **(B-C)** UMAP projection of 78,876 cells using 1,606 features per cell after removing ribosomal, mitochondrial, T-cell repertoire, and immunoglobulin genes. Cells are coloured by disease status **(B)** and broad cell type **(C)**. **(D)** Dot plot of the top three differentially expressed genes (DEGs) used to define PBMC broad clusters: T-cells, B-cells, natural killer (NK) cells, monocytes, dendritic cells, other myeloid cells, and hematopoietic stem cells (HSC). Clusters were annotated with SciBetR as well as manually curated. **(E)** Bar plot showing the cell abundance of broad clusters in PD and CTRL groups. Differential cell population abundance was computed using a beta-binomial generalized linear model. No significant percentage differences were observed: B-cells (*P =* 0.23), dendritic cells (*P =* 0.21), monocytes (*P =* 0.16), NK cells (*P =* 0.45), T-cells (*P* = 0.09). Error bars represent standard error of the mean. **(F)** DEGs in myeloid clusters: 99 in monocytes, 46 in dendritic cells, with 31 common genes (including: *CXCR4, ZFP36, RPS26, JUNB, FOS, CMEC12A, SAMHD1, FGL2, TYMP, RNF213*). (G) DEGs in lymphoid clusters: 44 in B-cells, 33 in NK cells, 65 in T-cells, with 10 common genes (*RPS26, JUNB, NFKBIA, RPS2, NR4A2, DUSP1, ZFP36, FOS, BTG1, ZFP36L2*).

Cell annotation was carried out using Seurat’s reference mapping procedure with datasets from Hao *et al*.^41^ and SciBetR v0.1.0, which utilized the COVID-19 PBMC Ncl-Cambridge-UCL reference dataset by Stephenson *et al*.^34^ Following this process, we observed comparable cell distributions between PD and CTRL within the UMAP space (Fig. 1B). We identified seven broad cell types, including T-cells, B-cells, NK cells, monocytes, DC, hematopoietic stem cells (HSC), and other myeloid cells (basophils, platelets), confirmed through manual curation based on the expression of classic marker genes (Fig. 1C, 1D). There were no significant differences in the overall proportions of the broad PBMC populations between the PD and CTRL groups. However, we observed trends showing lower percentages of T-cells (*P* = 0.09) and B-cells (*P =* 0.23), and higher percentages of monocytes (*P =* 0.16) and NK cells (*P =* 0.45) in the PD group (Fig. 1E). Our transcriptomic analysis between PD patients and healthy CTRL revealed significant differentially expressed genes (DEGs) across various immune cell types. Specifically, we identified 99 DEGs in monocytes, 46 in DC, 44 in B-cells, 33 in NK cells, and 65 in T-cells (Supplementary Fig. 2). When comparing DEGs between myeloid cells (monocytes and DC), 31 common DEGs were identified. Notable increases were observed in genes such as *CXCR4*, *ZFP36*, *RPS26*, *JUNB*, and *FOS*, while *CLEC12A*, *SAMHD1*, *FGL2*, *TYMP*, and *RNF213* showed decreased expression (Fig. 1F). Within the lymphoid cell compartment (T-cells, B-cells, and NK cells), 10 common DEGs were identified, all of which were upregulated in PD compared to CTRL. These genes include *RPS26*, *JUNB*, *NFKBIA*, *RPS2*, *NR4A2*, *DUSP1*, *ZFP36*, *FOS*, *BTG1*, and *ZFP36L2* (Fig. 1G).

This analysis revealed generally comparable proportions of the main immune cell types between PD patients and healthy controls. However, it also identified interesting DEGs across all immune cell types, indicating distinct regulatory patterns within the immune system in PD.

### Myeloid cells reveal a shift of classical monocytes to an activated phenotype in PD

Myeloid cells function primarily in innate immunity through phagocytosis, inflammatory response modulation, antigen presentation, and tissue repair.^45^ Through single-cell transcriptomic profiling, we identified various subpopulations within the myeloid lineage, which were categorized into specific DC and monocyte subsets (Fig. 2A). Initial clustering was performed using the SciBetR prediction algorithm, which identified 10 distinct clusters, including two separate CD14^+^ monocyte clusters and two CD16^+^ non-classical monocyte clusters. Manual curation of the CD14^+^ clusters revealed the presence of the activation marker CD83 in one cluster, which we annotated as CD14^+^/CD83^+^ or activated monocytes.^46,47^ Similarly, within the CD16^+^ clusters, we identified a cluster with high expression of the C1 complement genes, which we designated as CD16^+^/C1^+^ monocytes (Fig. 2B). These subsets also included AXL^+^SIGLEC6^+^DC (ASDC, transitional DC), pDC (plasmacytoid dendritic cell), DC1, DC2, DC3, proliferating DC (DC prolif) (Fig. 2A, 2B). Differential composition analysis using DCATS revealed significant differences between PD and CTRL groups. Specifically, we observed a significant decrease in the abundance of classical CD14^+^ monocytes (*P* = 0.03), accompanied by a concomitant increase of activated CD14^+^/CD83^+^ monocytes (*P* < 0.01) in PD patients (Fig. 2C), which was validated by multiple tools ( see Methods). Associated DEGs were extracted to investigate related molecular mechanisms, complemented by analysis of differences between PD and CTRL expression across all clusters (Supplementary Table 2 and 3). The ASDC population also showed a significant increase in abundance (*P* = 0.03). However, the cell count was extremely low (less than 1% of all cells), making further interpretation difficult.

**Figure 2.**
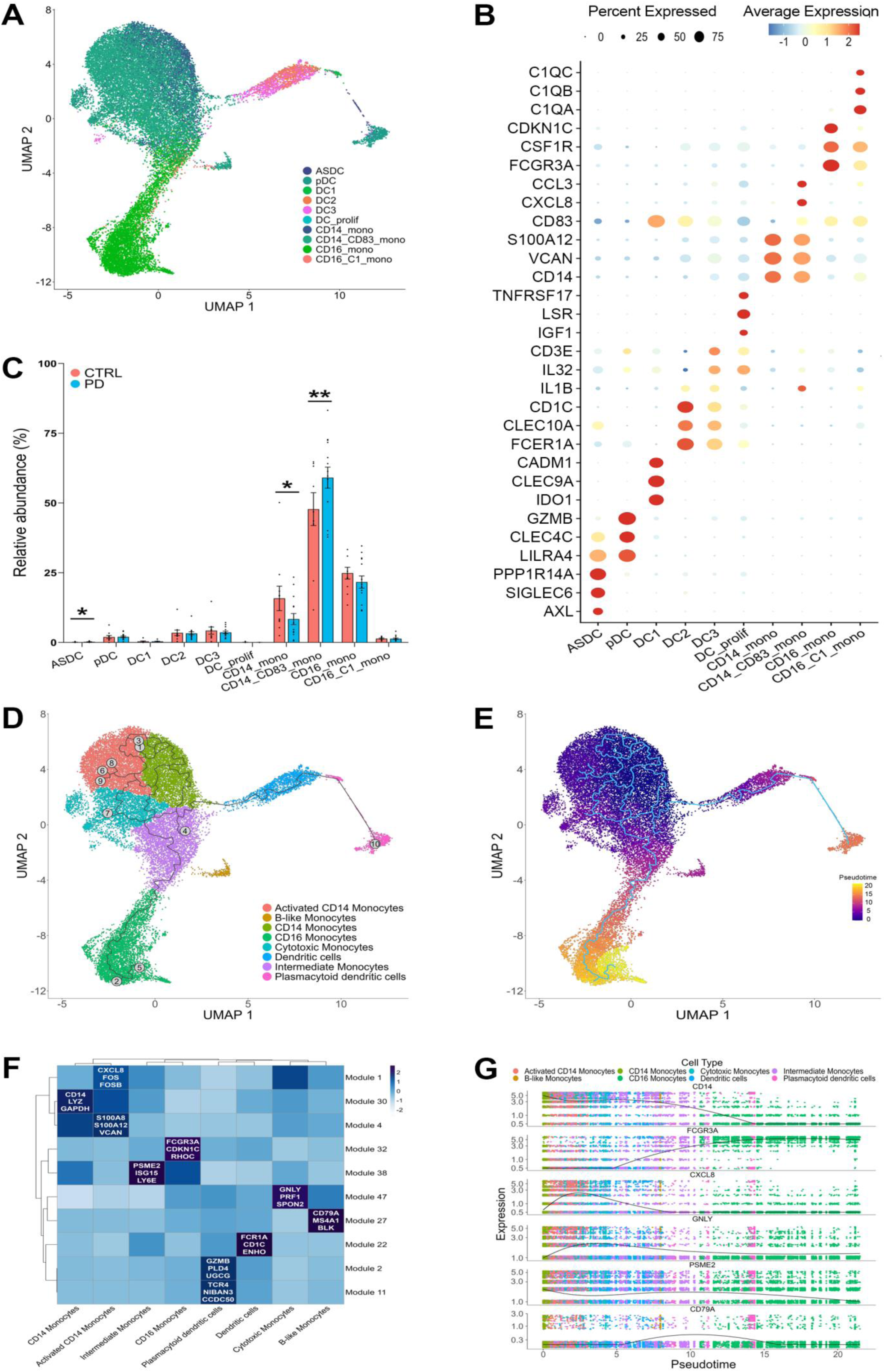
Single-cell transcriptomic profiling of myeloid cells revealed a shift of CD14^+^ classical monocytes to activated CD14^+^/CD83^+^ in PD. **(A)** UMAP projection of 18 169 cells from the myeloid population (monocytes, dendritic cells). Automated annotation followed by manual curation identified 10 subclusters; AXL^+^SIGLEC6^+^DC (ASDC), plasmacytoid DC (pDC), DC1, DC2, DC3, DC prolif, CD14^+^ monocytes, CD14^+^/CD83^+^ monocytes, CD16^+^ monocytes, CD16^+^/C1^+^ monocytes). **(B)** Dot plot of top three markers used to define the myeloid subclusters. Manual curation allowed the identification of the CD14^+^/CD83^+^ monocytes subcluster which are classical monocytes expressing the CD83 activation marker. The non-classical monocytes expressing the protein complement C1 (CD16^+^/C1^+^) were also identified through the same method. **(C)** Bar plot showing the cell abundance of myeloid cluster’s subpopulations in PD and CTRL. A significant difference was observed in CD14^+^/CD83^+^ monocytes with a higher abundance in PD (*P* < 0.01), with a concomitant lower abundance of CD14^+^ classical monocytes (*P* = 0.03). A significant increase in abundance was also observed for ASDC (*P* = 0.03) in PD. No significant difference was observed for pDC (*P =* 0.24), DC1 (*P =* 0.54), DC2 (*P =* 0.65), DC3 (*P =* 0.18), DC prolif (*P =* 0.50), CD16^+^ monocytes (*P =* 0.82), CD16^+^/C1^+^ monocytes (*P =* 0.59). Data were analyzed using DCATs beta-binomial generalized linear model. Error bars represent standard error of the mean. **(D)** Reclustering of the myeloid cells based on the same UMAP generated in (A) using Monocle3 pseudoclustering inference. A new cluster of cytotoxic monocytes was identified with this method. Numbers on the UMAP correspond to pseudotrajectory endpoints **(E)** Pseudotime analysis to identify the differentiation trajectories of the myeloid cells showing the evolution from classical monocytes to inflammatory states. **(F)** Overview of the main modules driving the cell differentiation. The top three genes in the most significant modules are highlighted. **(G)** Gene expression trajectory plot of top monocyte module genes along pseudotime.

Using Monocle 3, we re-clustered the myeloid cells based on DEGs, considering biological processes and alternative cell fates (Fig. 2D). This approach uncovered previously unrecognized subtypes, including a new cluster of cytotoxic monocytes and another characterized by B-cell signaling.^48^ This reclustering also refined the cells specifically associated to an activated monocyte trajectory (Fig. 2D). While statistical analysis of cell percentages did not reveal that specific subclusters were significantly driving the previously observed CD14^+^ monocytes differences between PD and CTRL (Fig. 2C), cell ratio trends support the idea that PD samples have less classical CD14^+^ monocytes (*P* = 0.05) while exhibiting a higher proportion of activated (*P* = 0.13) and cytotoxic monocytes (*P* = 0.13) (Supplementary Fig. 3).

Pseudotime analysis was conducted to map the differentiation trajectories of these cells and illustrated a progression from classical monocytes to more inflammatory profiles (Fig. 2E). To deduce cell identity, gene modules most associated with each refined cluster were compared with literature. The genes indicated common roles in specific immune pathways, such as cytotoxic functions (GNLY, PRF1, SPON2), B lymphocyte functions (CD79A, MS4A1, BLK), or cytokine response (ISG15, PSME2, LY6E). Additionally, several genes correlated with a stronger activation signal (CXCL8, FOS, FOSB, S100A8, S100A12, VCAN) in the remaining monocyte clusters (Fig. 2F). Gene expression trajectory for top markers from refined monocyte clusters illustrates how the genes transition across pseudotime (Fig. 2G).

Together, this analysis of myeloid cells revealed a shift of CD14^+^ classical monocytes to activated CD14^+^/CD83^+^ monocytes with diverging fates in PD, and highlights the dynamic changes in cell states and their potential roles in the inflammatory environment observed in PD.

### Transcriptomic profiling of lymphoid cell subpopulations

Lymphoid cells primarily function in adaptive immunity by producing antibodies, coordinating immune responses, direct cell killing through cytotoxic mechanisms, and generating immunological memory against specific pathogens.^50^ As with the myeloid lineage, we used SciBetR prediction followed by meticulous manual curation of cell markers to identify and categorize 24 distinct clusters within the lymphoid lineage (Fig. 3A; Supplementary Fig. 4). This approach allowed us to classify the three main lymphoid populations: T-cells (encompassing CD4^+^, CD8^+^, natural killer T cells (NKT), and mucosal-associated invariant T (MAIT) cells), B-cells, and NK cells, along with their respective subclusters.

**Figure 3.**
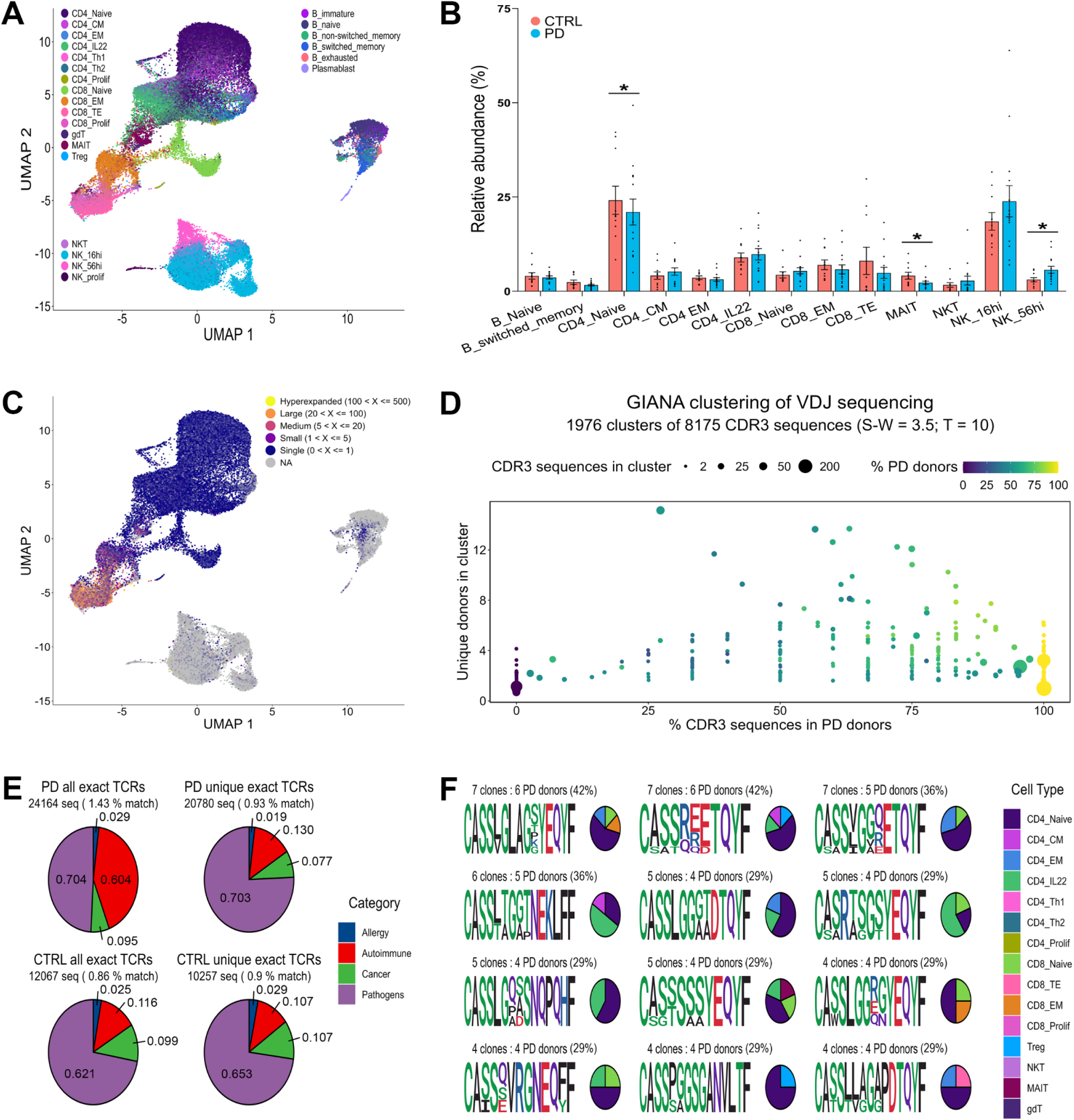
Transcriptomic profiling of lymphoid cell subpopulations. **(A)** UMAP projection of 52,954 cells from the lymphoid population (T-cells, B-cells, NK cells). Automated annotation followed by manual curation identified 24 subclusters (B immature, B naive, B non switched memory, B switched memory, B exhausted, plasmablast, CD4^+^ naive, CD4^+^ central memory (CM), CD4^+^ effector memory (EM), CD4^+^ IL22, CD4^+^ Th1, CD4^+^ Th2, CD4^+^ proliferating cells (CD4^+^ prolif), CD8^+^ naive, CD8^+^ EM, CD8^+^ T effector (TE), CD8^+^ proliferating cells (CD8^+^ prolif), gamma delta T (gdT), mucosal-associated invariant T (MAIT), regulatory T cells (Treg), NKT, NK16hi, NK56hi, NK proliferating cells (NK prolif)). **(B)** Bar plot showing the cell abundance of lymphoid cluster’s subpopulations in PD and CTRL. Subpopulations with a cell percentage below 1% relative to the lymphoid cluster were excluded from our analysis (B immature, B non switched memory, B exhausted, plasmablast, CD4^+^ Th1, CD4^+^ Th2, CD4^+^ prolif, CD8^+^ prolif, gdT, T reg, NK prolif). A significant difference was observed in CD4^+^ naive (*P* = 0.02) and MAIT (*P* < 0.01) which were less abundant in PD. Conversely, PD patients exhibited a significant increase in NK56hi (*P* = 0.03) abundance. No significant difference was observed for the other clusters: B naive (*P* = 0.11), B switched memory (*P =* 0.36), CD4^+^ CM (*P*= 0.74), CD4^+^ EM (*P =* 0.23), CD4^+^ IL22 (*P =* 0.89), CD8^+^ naive (*P =* 0.47), CD8^+^ EM (*P =* 0.14), CD8^+^ TE (*P =* 0.56), NKT (*P =* 0.15), NK16hi (*P =* 0.72). Data were analyzed using DCATs beta-binomial generalized linear model. Error bars represent standard error of the mean. **(C)** UMAP projection of clonal expansion observed from TCR sequencing. **(D)** Scatter plot of TCR sequence composition of clusters identified by GIANA v4.1 depicting cluster size, in number of CDR3 sequences (point size), percentage of sequences from PD donors (x axis), number of individual donors in cluster (y axis) and percentage of sequences coming from PD donors (point fill color). Yellow dots thus represent clusters containing CDR3 sequences only found in PD donors. 1,976 total clusters were identified. 364 clusters were PD specific, shared at least by two donors which correspond to 1 094 sequences, when excluding non-unique sequences. **(E)** Pie charts depicting predicted CDR3 sequence antigen specificity in CTRL vs PD cells by using the McPAS TCR database. Left column represents CDR3 sequences with exact matches in the database. Right column represents a unique exact method. Values expressed in percentage of total CDR3 sequences. **(F)** Peptide sequence logos and cell type composition for the top 12 TCR clusters with the most abundant shared PD-specific CDR3 sequences. Cell types of origin are illustrated along with the cluster sequences for top hits.

Differential composition analysis revealed decreases in CD4^+^ naive T-cells (*P =* 0.02) and MAIT cells (*P <* 0.01), as well as an enrichment of NK cells in PD, with a marked increase in the NK56hi (CD16^low^/CD56^++^) subpopulation (*P =* 0.03) (Fig. 3B). This subpopulation is known for its heightened immunoregulatory activity compared to the cytotoxic NK16hi (CD16^+^/CD56^low^), and its function suggest a potential role in the immune response alterations observed in PD.^51^

This analysis of the lymphoid cell compartments revealed decreases in subtypes of T-cells and increased cytotoxic NK cells. The increased presence of NK56hi cells in PD patients could indicate an enhanced state of immune activation or a compensatory mechanism in response to inflammation.

### TCR sequencing

T-cells play crucial roles in adaptive immunity by recognizing and killing infected cells, coordinating immune responses, and regulating the activity of other immune cells.^52^ Variable-diversity-joining (VDJ) recombination is a process of genetic rearrangement that occurs in developing T-cells and B-cells, which are critical components of the adaptive immune system. This process generates the diverse repertoire of antigen receptors necessary for recognizing a vast array of pathogens and are essential for a naive T-cell to undergo clonal expansion and differentiate into effector subsets. Therefore, analyzing the VDJ repertoire provides insight into the nature of an individual’s adaptive immune response. To explore the diversity and specificity of the T-cell population in PD patients, we performed single-cell sequencing of the variable regions of TCRs on the same cells that were analyzed for transcriptomic data and plotted the distribution of TCR expansion categories on the UMAP (Fig. 3C).

Identical TCR sequences indicate T-cell clonal expansion patterns and lineages, which are pivotal for recognizing endogenous and exogenous antigens presented by the major histocompatibility complex.^53^ In both PD and CTRL groups, the most expanded clones were predominantly found in the effector CD8^+^ T and NKT clusters, reflecting their high immune activity. However, the majority of clonotypes were detected within the CD4^+^ T-cell populations, owing to their higher abundance (Fig. 3C). Clustering of all CDR3 sequences based on similarity using the GIANA algorithm (see Methods) generated 1,976 clusters. Of those, 364 clusters representing 1,094 sequences were unique to PD cells and shared at least by two donors (Fig. 3D).^39^

We assessed if the identified CDR3 sequences had predicted specificity for antigens associated to disorders in the McPAS-TCR database.^54^ The “All exact” methodology is a clonal-expansion sensitive approach querying all CDR3 sequences for exact matches in the database on the overall avidity of our repertoire. The “Unique exact” results were obtained by removing duplicated sequences from a single donor, in order to assess the impact of single-patient clonal expansion (Fig. 3E). While exact repertoire coverage in the database was low at 0.86% and 1.43% for CTRL and PD respectively, it showed a striking enrichment of autoimmune-annotated CDR3 sequences in PD cells (Fig. 3E). The “Unique exact” profiles were similar between CTRL and PD groups (Fig. 3E), suggesting that the enrichment of the autoimmune signal came from clonally expanded sequences in specific PD patients. Detailed analysis revealed that the enrichment in autoimmune-annotated sequences in the “Unique exact” is driven by a single CDR3-sequence (CASSRDSNQPQHF) with a predicted reactivity to psoriatic arthritis and found in three distinct PD individuals, with a high clonotype expansion in one individual.

We next focused on epitope prediction of PD-specific CDR3 clusters shared between multiple donors. Interestingly, we identified clones shared by several PD patients, including two clones shared by 42.86% (6/14), two other clones shared by 35.71% (5/14) and ten clones shared by 28.57% (4/14) of the patients. However, none of the 364 shared clusters unique to multiple PD donors had predicted reactivity to antigens previously associated with PD (Fig. 3F). While the epitopes recognized by these PD-specific CDR3 sequence clusters are currently unknown, these results suggest common and shared PD-specific epitopes which could better inform us on PD pathogenesis.

These results reflect the diversity of CDR3 sequences in PD, spanning over 300 clusters shared among varying numbers of donors. More importantly, our results suggest a consistent immune mechanism among our PD patients, where a high percentage share common TCR clusters despite PD’s known heterogeneity.

### Increased activation signature gene expression in PD cell clusters

Our comparative DEG analysis revealed that several genes from the AP-1 transcription factor family as well as activation signature genes (ASG), recently described as stress response genes,^55,56^ were increased in both the myeloid and lymphoid cell populations in PD patients (Fig. 1F, 1G). Utilizing Seurat’s module scoring tool, we grouped the ASGs and visualized their expression levels across CTL and PD cell types (Fig. 4A). This revealed elevated ASG scores in PD vs CTRL across most cell types. To determine if the signal was an artifact of sample processing, we evaluated atypical parkinsonism participants (three MSA and three PSP) that were included at the beginning of our study (Fig. 4B). Intriguingly, their ASG scores were significantly lower than CTRL across the cell populations. Moreover, all the individual ASGs exhibited higher expression in PD compared to CTRLs, MSA and PSP (Fig. 4C). To identify which cell types contribute most to the ASG signal in PD, we examined the signal on the feature plot and the corresponding cell population. The ASG signal was the most elevated in CD14^+^/CD83^+^ activated monocytes, NK56hi cells, CD8^+^ EM T-cells and MAIT (Fig. 4D). For insights into the biological processes underlying the ASG signature, we performed pathway analysis on DEGs, focussing on differential cell types between CTL and PD; activated monocytes and NK56hi (Fig. 4E, 4F; Supplementary Fig. 5). A non-weighted KEGG analysis revealed a significant enrichment of two important immune pathways, IL-17 signaling pathway in activated monocytes (Fig. 4E), and NF-kB in both activated monocytes and in NK56hi (Fig. 4E, 4F). Furthermore, the NK56hi subpopulation exhibited enrichment in the TNF signaling pathway, another key proinflammatory pathway (Fig. 4F). Weighted GO term and KEGG pathway analysis also returned immune-relevant hits such as positive regulation of migration for CD83^+^ monocytes, innate immune response activation for NK56hi, and various neurodegenerative disorders for both cell populations (Supplementary Fig. 5).

**Figure 4.**
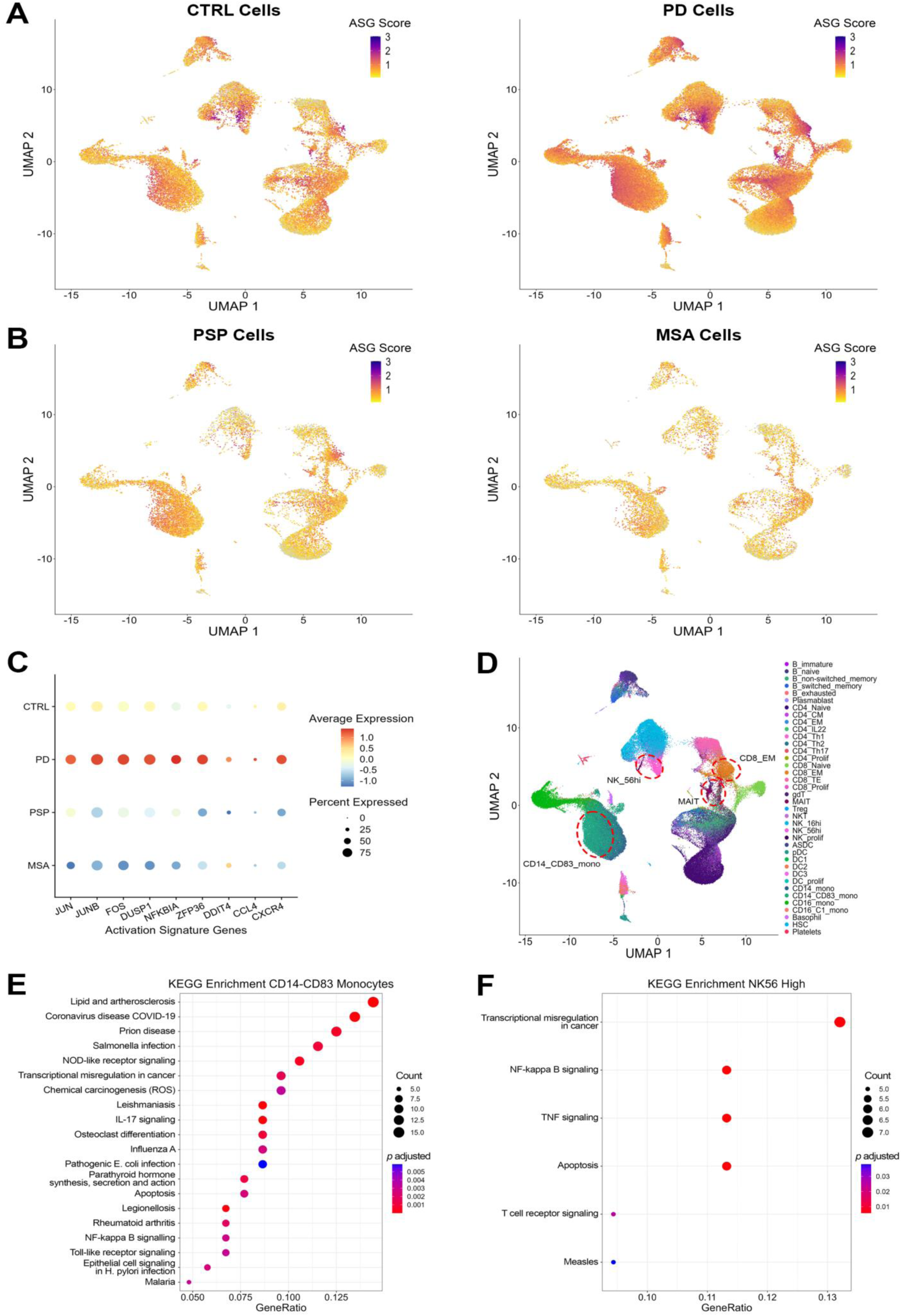
Activation signature genes. **(A)** Feature plot of ASG module score of all cells in PD and CTRL, showing an increased expression across most cell types in PD. **(B)** Feature plot of ASG module score of all cells in PSP and MSA. The expression of ASG genes based on the module score of these two phenotypes is lower compared to PD. **(C)** UMAP representing the ASG module hotspot (red dot lines), which are respectively the CD14^+^/CD83^+^ monocytes, the NK56hi, the CD8^+^ EM and the MAIT. **(D)** Dotplot showing the expression level of each ASG gene, in all phenotypes. In PD donors, most ASG genes have a *log2FC* > 1, while they are less expressed in CTRL, PSP and MSA. **(E-F)** Non-weighted KEGG analysis of DEGs showing the top 20 most significant pathways in CD14^+^/CD83^+^ activated monocytes **(E)** and NK56hi **(F)**. Activated monocytes show enrichment in immune-related pathways (e.g IL17 signaling pathway, NF-kappa B signaling pathway), in metabolism (lipid and atherosclerosis), and in autoimmunity (rheumatoid arthritis) **(E)**. NK56hi pathway analysis also revealed immune-related pathways (NF-kappa B signaling pathway, TNF signaling pathway, TCR signaling pathway) **(F)**.

Together, these data reveal consistent dysregulation of ASGs across both myeloid and lymphoid cells in PD, but not in atypical parkinsonism, markedly in over-represented populations such as activated monocytes (Fig. 2C) and NK56hi cells (Fig. 3B).

## Discussion

### From broad clusters to specific subpopulations

Single-cell transcriptomics was used in a funnel-down manner to analyze the complete PBMC compartments simultaneously, initially focusing on broad clusters and progressively honing in on specific populations.

While the broad analysis found no significant difference in relative cell abundance between PD and CTRL broad PBMC clusters, PD samples trended towards an enrichment in monocytes and a reduction in lymphocytes (Fig. 1E). Although none of the previous sc-RNAseq publications encompassed the entire PBMC compartment in PD, Tian *et al*.^57^ reported similar findings, showing significant increases only in the ratio of PD patients monocytes using flow cytometry analysis on total PBMC samples. Harnessing the power of single-cell analysis, we refined our annotations of myeloid and lymphoid compartments to deepen our analysis of each cell subtype and identify the subcluster driving those differences.

### An activated state of monocytes in PD

Marker based identification of the monocyte population allowed us to identify a subcluster of CD14^+^ monocytes marked by the CD83 immune-cell activation marker (Fig. 2B).^46^ CD83^+^ monocytes DEGs include several chemokine signaling genes such as *CCL3*, *CXCL8*, *IL1B*, and *CXCR4* (Supplementary Table 2)*. CXCR4* overexpression has been previously described in the substantia nigra in PD and is associated with microglial activation. ^58^ Therfore, we hypothesize the observed CXCR4 overexpression in the periphery could contribute to the activation of peripheral immune cells, specifically monocytes. This potential involvement of CXCR4 in PD suggests that other related chemokines, some of which are differentially expressed in PD as mentioned earlier, might also be implicated.^59^ Consequently, these chemokine related signaling pathways may contribute to monocyte activation.

Classical CD14^+^ monocytes can be triggered to express CD83 when treated with the cytokine IFN-α. These activated monocytes exhibit phagocytic behavior similar to CD14^+^ monocytes and do not adopt a dendritic morphology, distinguishing them from DCs.^60^ IFN-α is a key cytokine in the response to viral infections, initiating the innate immune response. Under physiological conditions, lower levels of IFN-α/β are present without viral infections, resulting in a weak signal that primes cells for an amplified response during viral infections and increases their sensitivity to other cytokines.^61^ Based on our findings, it is possible that PD patients may have elevated baseline levels of circulating IFN-α, which could lead to a higher abundance of activated monocytes. Furthermore, Cao *et al*.^62^ demonstrated the presence of preformed CD83 within monocytes using Western blotting and flow cytometry, and showed that CD83 could be induced on the surface of monocytes following LPS stimulation, with expression peaking 2-4 hours post-stimulation and disappearing after 24 hours. This indicates that CD83 is not constitutively expressed but can be rapidly upregulated in response to specific stimuli. These observations suggest that PD patients might exhibit heightened sensitivity to stimulation and stress compared to healthy controls, resulting in a higher expression of *CD83* in CD14^+^ monocytes.

Consistent with these findings, our non-weighted KEGG pathway analysis in the activated monocyte populations revealed enrichment in multiple immune-related processes including certain previously linked to PD such as IL17 and NF-kB (*P* < 0.05) (Fig. 4E). IL-17 is a proinflammatory cytokine involved in chronic inflammation and present in the pathogenesis of autoimmune diseases such as rheumatoid arthritis, which generated a noticeable signal among our PD TCR sequences (Fig. 3E).^63,64,65,66^ Consistent with this, an increase of IL-17 producing cells as well as concentration in PD patient serum was reported by Yang et al. ^67^ Intriguingly, the proinflammatory NF-kB signaling pathway, which also emerged from the KEGG analysis, is activated by the IL-17 cytokine.^68^

Together, our results indicate that the activated state of CD14^+^/CD83^+^ monocytes may reflect baseline immune activation or immune dysregulation in PD. In addition, our findings provide additional evidence for the importance of inflammation and autoimmunity in PD pathogenesis, highlighting the specific role CD14^+^/CD83^+^ activated monocytes may play.

### Higher activation of NK56hi cells in PD

NK cells represent the innate immune system lymphocytes. Besides their cytotoxic and cytokine producing functions, they have an immunoregulatory role through their interaction with other immune cells.^69^ Here, we found a significant increase of regulatory NK56hi cells in PD compared to CTRL (*P =* 0.01) (Fig. 3B). Similar results were described in a flow cytometry study assessing the expression of the activating receptor NKG2D on NK cells between CTRL and PD subgroups.^70^ In this study, they used the Unified Parkinson’s Disease Rating Scale (UPDRS) to stratify a mild symptoms group and moderate to severe symptoms group.^71^ Interestingly, they found an increased NKG2D mean fluorescence intensity on NK56hi cells of PD patients only in the moderate/severe group.^70^

Accordingly, future sc-RNAseq investigations should attempt to analyse results following segregation by disease severity. However, an increased number of participants will be needed to attain sufficient statistical power.

The significant enrichment of GO terms relating to immune activation in our pathway analysis, along with the increased cell abundance, suggests a state of hyperactivation of the NK56hi cells, possibly in response to an immune dysregulation. Noticeably, our KEGG analysis identified significant enrichment of the NF-kB and TNF-α pathways (Fig. 4F). TNF-α is a pro-inflammatory cytokine which mostly leads to cell death, and its neutralization is utilized in chronic inflammatory and autoimmune disease treatment.^72^ Interestingly, NK cells produce TNF-α,^73^ which is intertwined with NF-κB as the latter is activated by TNF in most canonical NF-κB signalling pathways.^74,75^

Overall, this suggests a specific increase in activation of the immunoregulatory NK56hi in PD, which supports the hypothesis of immune activation or dysregulation in patients.

### Common TCR in PD

Recently, the notion of PD as a single entity has been challenged in favor of a more heterogeneous disease.^76^ Despite this heterogeneity, we uncovered 1,094 PD-specific sequences including several TCR clonotypes shared by up to 42% of our cohort, supporting the involvement of peripheral immune mechanisms irrespective of genetic backgrounds and environmental factors. Examining annotations associated with the GIANA “all exact” clusters showed that the TCR sequences in PD patients were enriched for “autoimmunity” (Fig. 3E). Indeed, three patients shared a sequence, which was expanded in only one of them, associated with psoriatic arthritis. None of these patients exhibited symptoms of psoriatic arthritis at the time of enrollment. Of note, databases such as McPAS-TCR still contain a limited amount of data. Thus, the low cluster match score (∼1%) suggests a high prevalence of CDR3 sequences with unidentified related pathology (Fig. 3E), highlighting the need to enrich annotation databases.

Wang *et al*.^20^ reported a similar identification of 67 PD-specific TCR groups. Using machine learning, they screened these TCRs and their candidate epitopes to discover that predicted antigens could induce helper and cytotoxic T-cell responses in PD patients. Further analysis of our shared PD epitopes with emerging algorithms of this rapidly evolving field, coupled with future referencing with HLA-peptide/TCR experiments, could enable deeper insights into what triggers common immune responses in PD patients.^77,78^

### T-cell population imbalance

Sc-RNAseq studies in PD PBMCs have been limited to preselected subpopulations. Wang *et al.*^20,21^ conducted two single-cell studies, focusing separately on T-cell populations with TCR-sequencing, and on B-cells with BCR-sequencing.

Their T-cell analysis found increased CD4^+^ and CD8^+^ populations in PD samples. However, when looking at the complete lymphoid compartment, we observe a significant decrease of CD4^+^ naive cells in our PD cohort and no significant differences in the CD8^+^ population.^21^ Similarly, we did not find a differential abundance in our B-cell populations. In contrast, Wang *et al.*^21^ reported a significant increase of unswitched memory B-cells, along with a significant decrease of naive B-cells.

The discrepancy between Wang et *al.* results and ours may be attributable to differences in methodology such as the inclusion of pre-processing selection steps which may limit the reliability of quantitative population comparisons.

### Activation Signature Genes

In a 2022 publication, Marsh *et al.*^55^ identified a signal in their single-cell transcriptomics of PBMCs, which seemingly correlated to their tissue processing method. They showed that if you add a 30-minute enzymatic digestion at 37°C to a standard PBMC isolation protocol, cells upregulate several stress-response genes such as *JUNB, JUN, FOS, DUSP1, NFKBIA, ZFP36, DDIT4, CXCR4*, and *CCL4*, which they termed ASGs. The article recommends using transcription and translation inhibitors (TTIs) immediately upon blood draw to mitigate these “artifacts”. Subsequently, we found that ASGs were increased by standard PBMC isolation protocols, even without enzymatic digestion, and that adding TTIs or keeping cells cold (4°C) during processing could abrogate this signal.^56^ Moreover, we demonstrated that cells treated with TTIs no longer respond to immune stimulation and thus keeping cells cold would be more advantageous due to its flexibility and cost-effectiveness.

Surprisingly, while we observed increased ASG signal in all cells, it was significantly higher in PD samples. Importantly, when compared to other atypical parkinsonism donors processed at the same time, this increase was not present ; instead, lower ASG levels than CTRL samples were observed in both atypical parkinsonism groups (Fig. 4A, 4B, 4C). Despite the global overexpression of ASGs across PD-cell populations, the signal peaks correlate to previously described activated monocytes and NK56hi clusters (Fig. 4D).

Interestingly, recent studies have linked ASGs and AP-1 complex genes in particular, to aging immune-cell phenotypes. Karakaslar *et al.*^79^ described an epigenetically modulated age-related signature involving the AP-1 complex genes *JUN*, *JUNB*, and *FOS*. Human blood ATAC-sequencing revealed increased binding sites for these genes with age, potentially triggering the production of proinflammatory IL6 and contributing to the inflammaging process.^80^ This raises questions about the pathophysiology of immune dysregulation in PD, whether it is due to an impairment in the physiological process of inflammaging, or a reaction due to other disease-related mechanisms.

It is important to emphasize that all the samples in our study were processed using identical methods, and each processing batch contained samples from all phenotypes, so it is unlikely that processing artifacts led to this differential ASG signal. This reinforces the hypothesis of a heightened state of activation in PD. What remains to be determined is whether a higher baseline signal can be observed *in vivo* or if differential levels arise from stress responses during PBMC isolation and/or single-cell transcriptomic processing.

Overall, this intricate molecular and cellular signature in peripheral tissues highlights systemic immune dysregulation in PD patients, suggesting that targeting this hyperreactive immune state could be a promising therapeutic strategy. The comprehensive PBMC landscape atlas presented in this study aims to serve as a valuable resource for researchers, enhancing our grasp of the disease and facilitating the identification and validation of potential biomarkers and drug candidates.

## Acknowledgements

The authors are grateful to the patients and their families for their participation in this study. We would also like to thank the André-Barbeau movement disorders unit for blood sampling within the framework of a biobank, as well as the sequencing platform of the CHU de Québec-Université Laval’s genomic center. This research was enabled in part by the bioinformatic resources obtained through the Digital Research Alliance of Canada and the support of Calcul Quebec.

## Author contributions

Conceptualisation: G.M-B, M.T; Software: G.M-B, S.A; Formal Analysis: G.M-B, L.A, S.A, L.K.H; Investigation: G.M-B, L.A; Resources: A.D, S.C, M.P, M.T; Data Curation: S.A; Writing original draft: G.M-B, L.A, L.K.H; Writing Review & Editing: S.A, L.A, L.K.H, A.D, S.C, M.P, M.T; Visualization: G.M-B, S.A; Supervision: L.K.H, M.T; Funding acquisition: M.T

