## Supplementary figures and images for "Mapping the peripheral immune landscape of Parkinson’s disease patients with single-cell sequencing"

### Supplementary Fig. 1

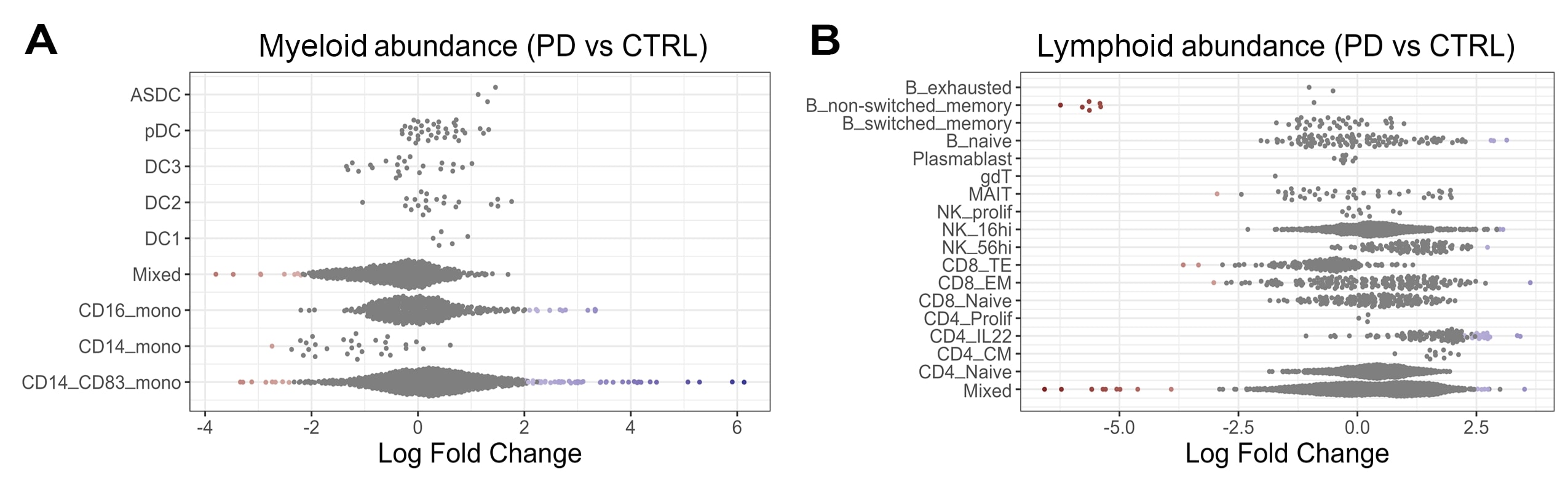

### Supplementary Fig. 2

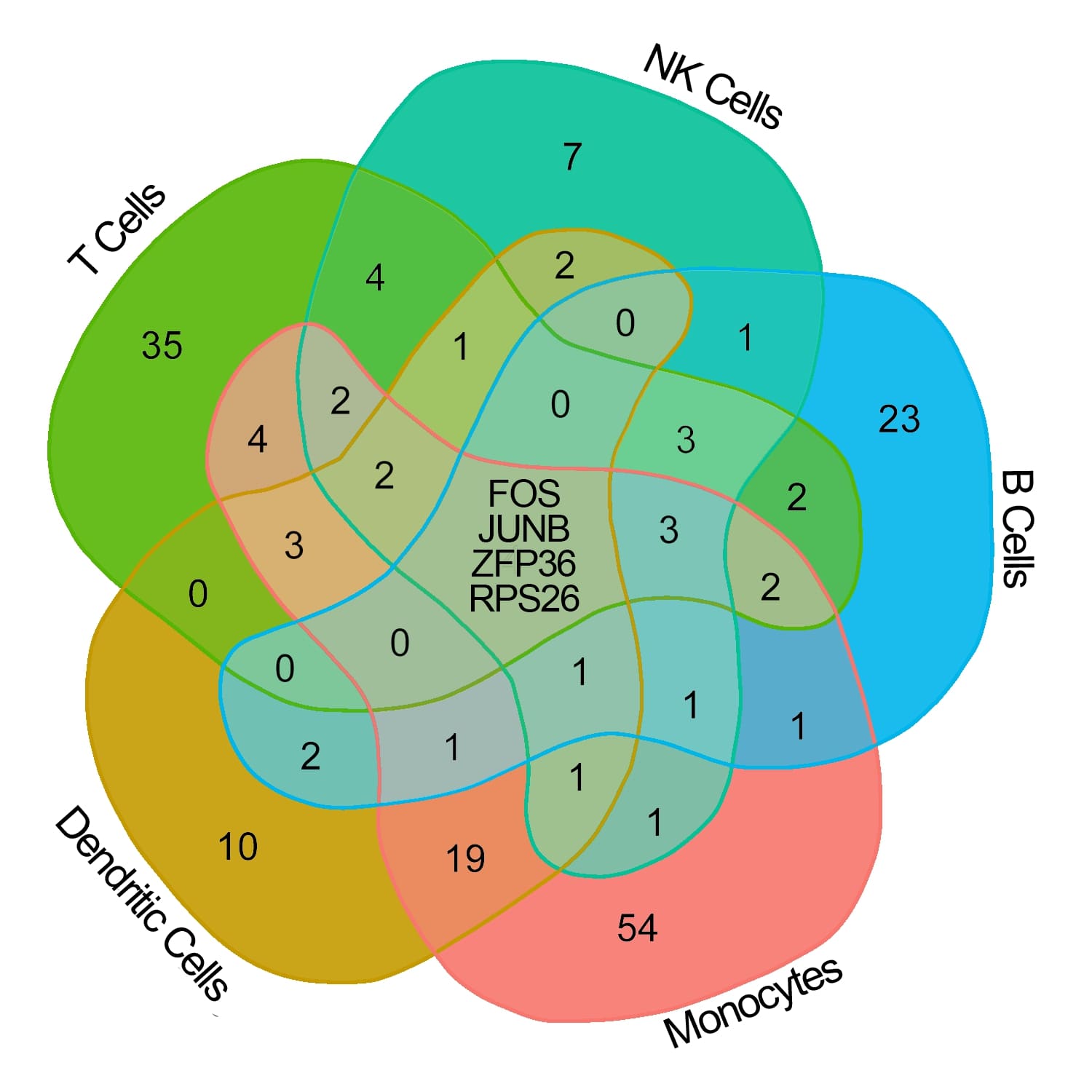

### Supplementary Fig. 3

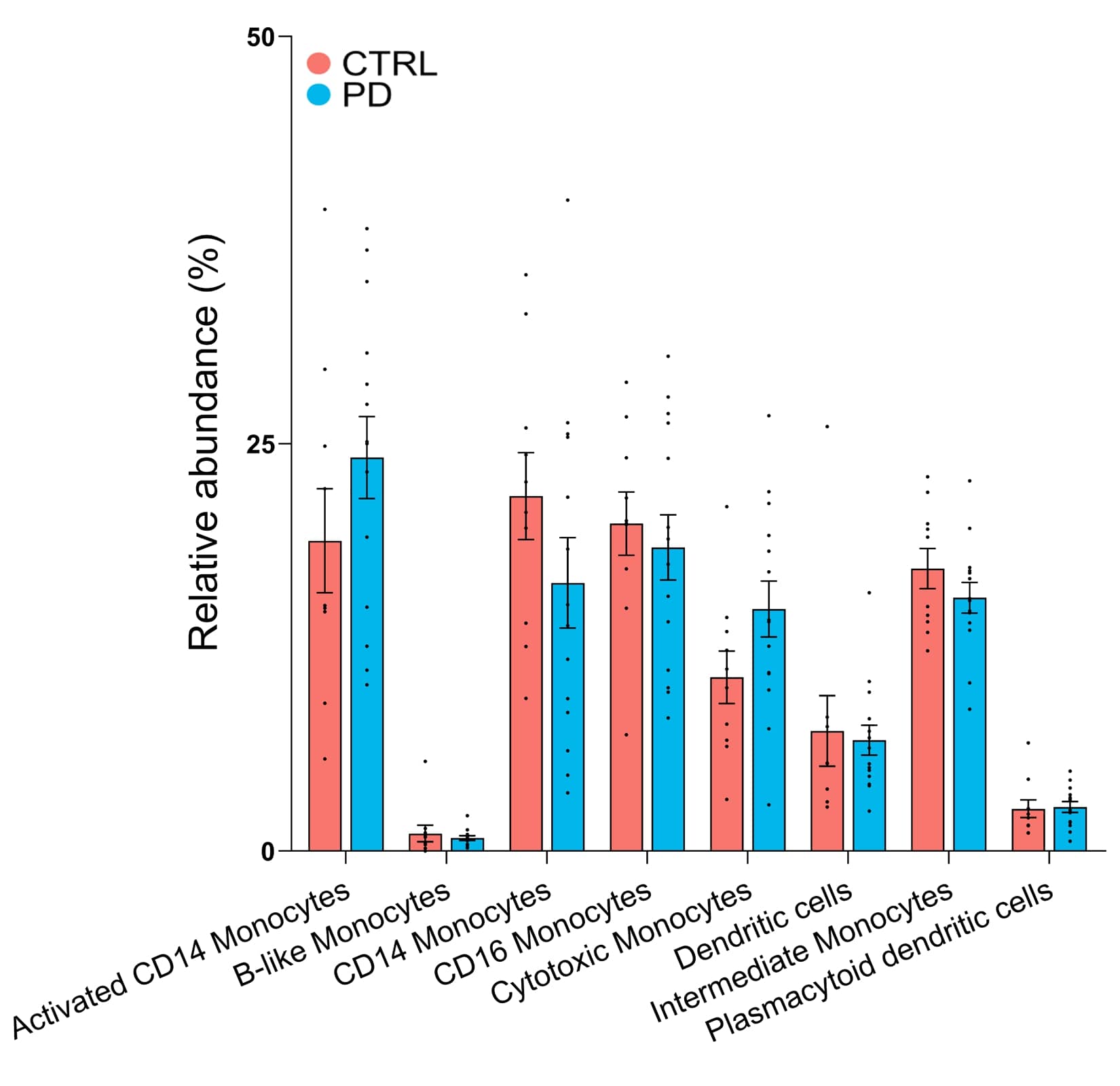

### Supplementary Fig. 4

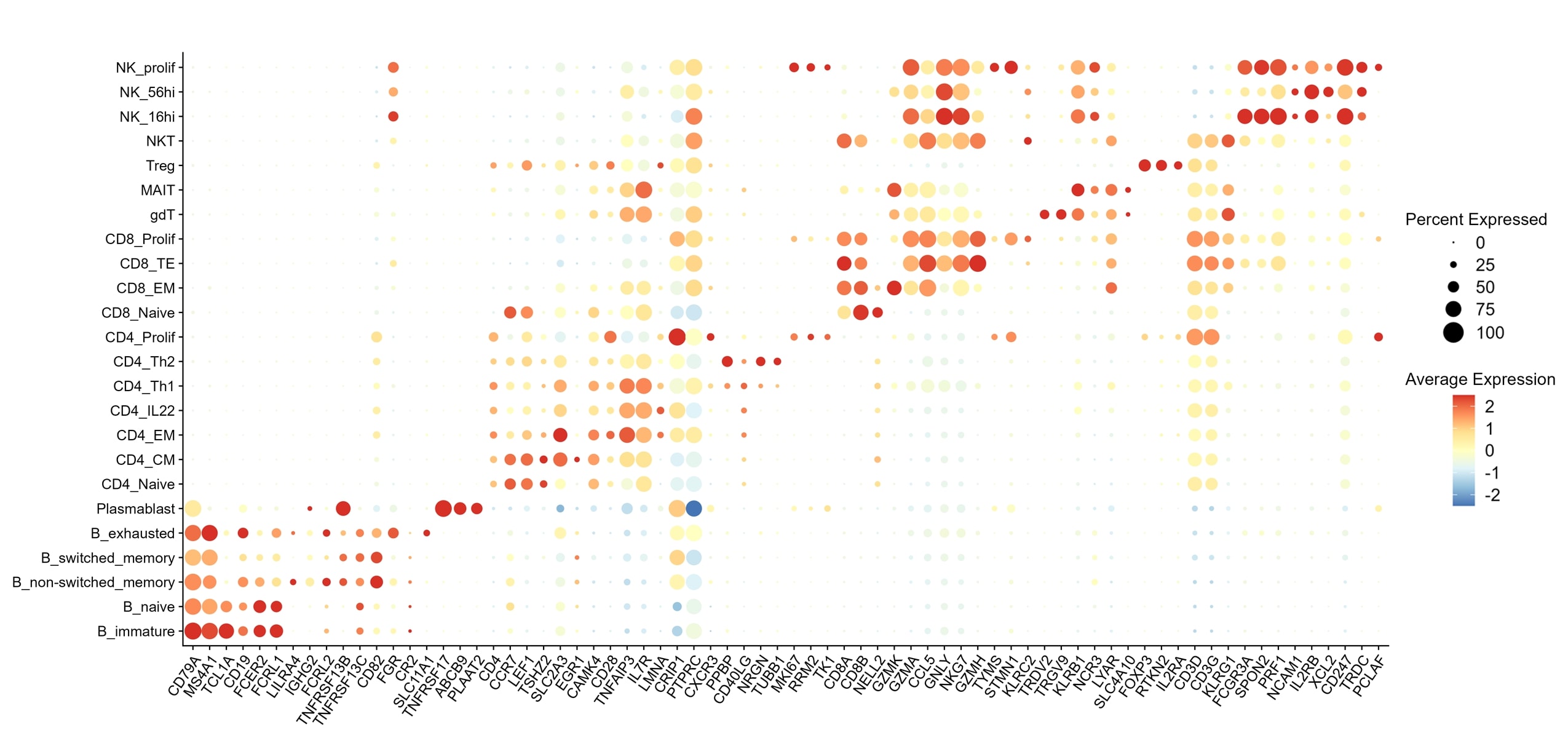

### Supplementary Fig. 5

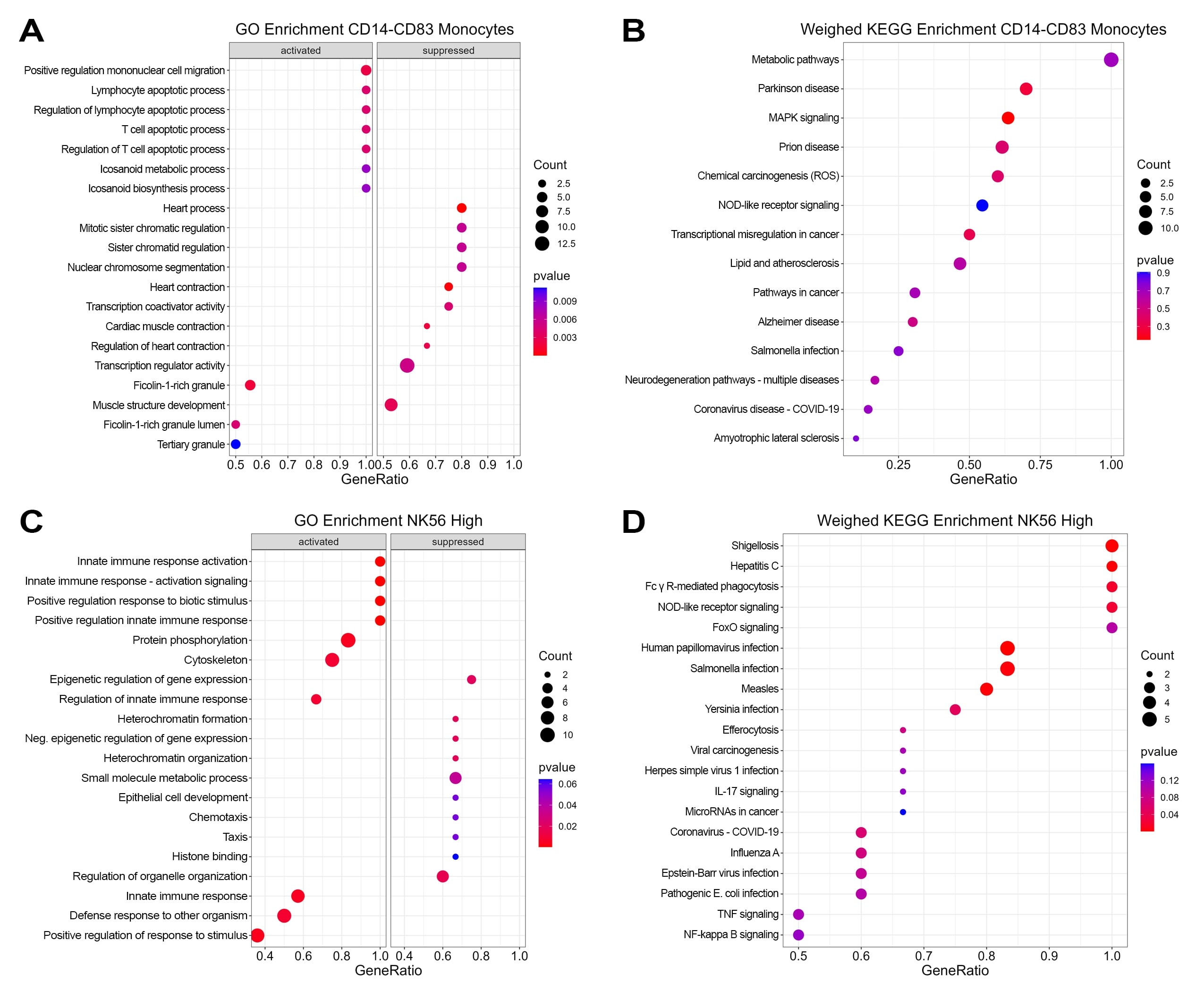
