## Supplementary Table 1 for "Mapping the peripheral immune landscape of Parkinson’s disease patients with single-cell sequencing"

| ID | Patient or CTRL | Diagnosis | Sex | Age | Age at onset | Duration of symptoms | Total UPDRS score |
|---|---|---|---|---|---|---|---|
| Prat-2020-10 | CTL | | F | 82 | | | - |
| Prat-2020-14 | CTL | | M | 80 | | | - |
| Prat-2020-08 | CTL | | M | 67 | | | - |
| Prat-2020-06 | CTL | | M | 63 | | | - |
| DM006C | CTL | | F | 72 | | | - |
| UTMAB-2020-0004 | CTL | | F | 64 | | | - |
| UTMAB-2021-0046 | CTL | | F | 66 | | | - |
| UTMAB-2021-0048 | CTL | | F | 66 | | | - |
| DM016C | CTL | | M | 67 | | | - |
| DM009C | CTL | | M | 52 | | | - |
| DM024P | Patient | MSA | M | 63 | 58 | 5 | 78 |
| DM027P | Patient | MSA | M | 64 | 63 | 2 | 40 |
| DM030P | Patient | MSA | M | 65 | 58 | 7 | 108 |
| UTMAB-2021-0043 | Patient | PD | M | 72 | 66 | 6 | 61 |
| UTMAB-2021-0047 | Patient | PD | M | 67 | 46 | 21 | 106 |
| DM029P | Patient | PD | M | 65 | 56 | 6 | 75 |
| UTMAB-2021-0045 | Patient | PD | M | 67 | 58 | 9 | 42 |
| UTMAB-2021-0056 | Patient | PD | F | 74 | 63 | 11 | 50 |
| UTMAB-2021-0040 | Patient | PD | F | 69 | 67 | 2 | 57 |
| DM008P | Patient | PD | M | 74 | 59 | 15 | N/A |
| UTMAB-2021-0049 | Patient | PD | M | 71 | 64 | 7 | 100 |
| DM007P | Patient | PD | F | 70 | 65 | 5 | N/A |
| UTMAB-2021-0041 | Patient | PD | F | 64 | 59 | 5 | 45 |
| UTMAB-2021-0044 | Patient | PD | F | 62 | 56 | 6 | 31 |
| UTMAB-2021-0057 | Patient | PD | M | 64 | 58 | 6 | 46 |
| UTMAB-2021-0039 | Patient | PD | F | 68 | 57 | 11 | 64 |
| DM023P | Patient | PD | F | 62 | 59 | 3 | 34 |
| Prat-2020-05 | Patient | PSP | F | 63 | 59 | 4 | 121 |
| Prat-2020-11 | Patient | PSP | M | 66 | 61 | 5 | 126 |
| Prat-2020-13 | Patient | PSP | F | 73 | 69 | 4 | 120 |

**Supplementary table 1. Cohort Demography.** Our single-cell study included 10 healthy controls and 14 patients with PD. The patients and controls were age- and sex-matched, although there were fewer controls. Additionally, we recruited patients with atypical parkinsonism, including three with MSA and three with PSP. (N/A = not available)
